# Integrating CRISPRi perturbation responses and transcription-factor binding priors for regulatory network inference in triple-negative breast cancer

**DOI:** 10.64898/2026.11.20.768082

**Authors:** Omar Faroque, Mst. Homayra Jannaty, Shahriar Hasan Abir, Suman Biswas

## Abstract

Triple-Negative Breast Cancer (TNBC) represents an aggressive breast cancer subtype that lacks actionable cell-surface receptors, restricting frontline systemic management to cytotoxic chemotherapy. While recent functional epigenomics shows that TNBC progression is governed by non-coding regulatory remodeling at distal super-enhancers, conventional Gene Regulatory Network (GRN) inference methods rely on observational transcriptomic co-expression, which suffers from technical dropouts and confounding correlations that inflate false discoveries (>30%). Here, we present an interventional network inference framework that integrates large-scale single-cell CRISPR-interference perturbation sequencing (Perturb-seq; GSE224986, 95,678 cells across 11,975 sgRNAs) with ENCODE Factorbook ChIP-seq physical transcription factor binding priors across 819 somatic mutation-associated enhancer loci in human TNBC (MDA-MB-231). Our pipeline reconstructed a tripartite network yielding 5,935 physical TF–enhancer edges, 383,757 functional enhancer–gene edges, and 542,760 unique TF–target gene functional pairs across 372 active enhancers. Topological centrality prioritized master transcriptional drivers governing TNBC oncogenic signaling, including CTCF, POLR2A, EP300, JUND, MYC, and FOS, which regulate Hallmark programs for Epithelial-Mesenchymal Transition (p = 1.2 × 10⁻⁹) and cell cycle progression. Explainable machine learning (SHAP TreeExplainer) revealed non-linear, thresholded transactivation dynamics for AP-1 and MYC. Quantitative benchmarking against held-out CRISPRi ground truth demonstrated strong discriminatory accuracy (AUROC = 0.8999, AUPRC = 0.8745, Precision = 90.13%, FDR = 9.87%), substantially outperforming GENIE3 (AUPRC = 0.6323, FDR = 35.73%) and co-expression correlation (AUPRC = 0.6814). In silico-simulated driver feature silencing correlated with experimental responses (R² = 0.473, p < 10⁻¹²²), and master driver signatures significantly stratified overall survival in the METABRIC (n = 320) and TCGA-BRCA (n = 160; p ≤ 0.019) cohorts. Finally, in silico perturbation modeling indicated that multi-target epigenetic combinations achieved 93% topological disruption of oncogenic regulatory connectivity, nominating prioritized candidate circuits for experimental validation.

**Author Summary:** Triple-negative breast cancer (TNBC) is one of the most challenging malignancies to treat because cancer cells lack the hormonal and HER2 receptors targeted by standard precision drugs. While tumor cells maintain their aggressive behavior by rewiring genetic control switches through specialized regulatory DNA regions known as super-enhancers, traditional computer models struggle to map these networks accurately because they rely on simple correlation, which often mistakes coincidental patterns for genuine biological cause and effect.

In this study, we addressed this limitation by developing a computational framework that directly links large-scale single-cell CRISPR-based genetic silencing experiments to physical protein-DNA binding maps. By analyzing how systematic genetic perturbations affect gene activity across nearly 100,000 individual cancer cells, our model identified master regulatory proteins, such as AP-1, MYC, and EP300, that drive tumor progression and correlate with poor patient survival in clinical cohorts. Furthermore, our computational simulations demonstrate that simultaneously targeting multiple epigenetic switches disrupts cancer networks far more effectively than standard chemotherapy, providing concrete, prioritized targets for future laboratory experiments and combination drug development.

## Introduction

### Clinical Gravity, Heterogeneity, and Therapeutic Challenges of TNBC

Triple-Negative Breast Cancer (TNBC) accounts for approximately 15% to 20% of all diagnosed breast carcinomas and remains disproportionately responsible for breast cancer mortality worldwide (Dent et al., 2007; Foulkes et al., 2010; Zagami & Carey, 2022). Clinically defined by the immunohistochemical absence of estrogen receptor (ER) and progesterone receptor (PR) expression, together with the absence of human epidermal growth factor receptor 2 (HER2) overexpression or gene amplification (Irvin & Carey, 2008; Rakha et al., 2007), TNBC represents a biological class that is fundamentally unresponsive to endocrine modulators (such as tamoxifen and aromatase inhibitors) or HER2-directed monoclonal antibodies (such as trastuzumab) (Dass et al., 2021; Waks & Winer, 2019). Consequently, systemic cytotoxic chemotherapy, comprising anthracyclines, taxanes, and platinum-based DNA-crosslinking agents, remains the historical mainstay of systemic treatment (Bianchini et al., 2016; Waks & Winer, 2019).

Despite aggressive multi-agent chemotherapy regimens, patient clinical trajectories are characterized by high relapse rates. While initial chemosensitivity is observed in a subset of patients, over 30% to 40% experience rapid disease recurrence within three to five years following definitive surgery (Dent et al., 2007; Foulkes et al., 2010). Metastatic TNBC exhibits an aggressive natural history with a pronounced predilection for visceral organ dissemination, including the lungs, liver, and central nervous system, resulting in a median overall survival of only 12 to 18 months (Bianchini et al., 2016; Garrido-Castro et al., 2019a). Comprehensive transcriptomic profiling has established that TNBC is an intensely heterogeneous disease encompassing distinct molecular entities, predominantly the basal-like (BL1, BL2), mesenchymal (M), mesenchymal stem-like (MSL), immunomodulatory (IM), and luminal androgen receptor (LAR) subtypes (Burstein et al., 2015; Lehmann et al., 2011; Perou et al., 2000; Sørlie et al., 2001). Because standard targeted kinase or cell-surface receptor inhibitors consistently fail due to rapid bypass signaling and secondary resistance (Garrido-Castro et al., 2019a), there is an urgent requirement to unravel the core transcriptional and epigenetic regulatory circuits that sustain TNBC survival and metastatic fitness.

### Epigenetic Architecture, Super-Enhancers, and Non-Coding Remodeling

Recent advances in cancer functional genomics demonstrate that oncogenic states are not solely governed by focal protein-coding mutations, but are intensely dependent on non-coding regulatory elements, particularly enhancers and super-enhancers(Bradner et al., 2017; Hnisz et al., 2013; Lovén et al., 2013; Whyte et al., 2013). Super-enhancers are dense clusters of broad genomic enhancers characterized by exceptional occupancy of Mediator (MED1), the histone acetyltransferase EP300/CBP, and high enrichment of histone H3 lysine 27 acetylation (H3K27ac) (Whyte et al., 2013). Through phase-separated biomolecular condensation and long-range chromatin looping mediated by the CCCTC-binding factor (CTCF) and cohesin complexes (Sabari et al., 2018; Zhao et al., 2021), super-enhancers compartmentalize the transcription apparatus to drive exceptionally high-level expression of lineage-defining master regulators and oncogenes (Bradner et al., 2017; Lovén et al., 2013).

In basal-like TNBC, malignant cells undergo extensive epigenetic remodeling that establishes de novo super-enhancers at loci driving proliferation, stemness, and epithelial-to-mesenchymal transition (EMT) (Franco et al., 2018; Lewis et al., 2024). Active enhancers frequently undergo bidirectional transcription into non-coding enhancer RNAs (eRNAs), which stabilize chromatin loops and facilitate RNA polymerase II elongation (Li et al., 2016). In recent functional screening investigations, (Lewis et al., 2024) and (Wang et al., 2025) deployed pooled single-cell CRISPR-interference (CRISPRi) tiling across transcribed super-enhancers and non-coding somatic mutation enhancer loci in the archetypal TNBC line MDA-MB-231, demonstrating that targeted silencing of specific distal super-enhancers severely impeded tumor growth and metastatic progression in vivo. Nevertheless, despite the identification of individual functional enhancers, resolving the complete directed regulatory network (identifying which transcription factors physically occupy these elements and which downstream target genes they causally govern) has remained an elusive bioinformatic and statistical challenge.

### Inferential Challenges of Observational GRN Methods and Opportunities from Single-Cell CRISPRi

Historically, Gene Regulatory Networks (GRNs) have been inferred from observational transcriptomic datasets using correlation metrics (Pearson, Spearman), mutual information algorithms (ARACNE, CLR) (Faith et al., 2007; Margolin et al., 2006), or non-linear regression ensembles (GENIE3, SCENIC) (Aibar et al., 2017; Huynh-Thu et al., 2010). However, the systematic application of observational algorithms to single-cell RNA-sequencing (scRNA-seq) has exposed catastrophic methodological limitations (Chan et al., 2017; S. Chen & Mar, 2018; Pratapa et al., 2020). Single-cell data suffer from severe technical zero-inflation, commonly designated as “dropout,” arising from low mRNA capture efficiency and stochastic amplification bias (Dai et al., 2024). In that study, the authors provided rigorous mathematical proof that dropout zeros function as systematic confounders that alter joint probability distributions, inducing artificial statistical dependencies between completely unrelated genes.

Furthermore, observational correlation cannot distinguish direct causal regulation from transitive dependencies or shared upstream confounding (Pearl, 2009; Peters et al., 2017; Spirtes et al., 2001). For instance, if master regulator A transcriptionally activates both intermediate gene B and target gene C, genes B and C will exhibit strong observational co-expression despite having zero regulatory relationship between them. In rigorous benchmarks against biological ground truth, observational GRN algorithms routinely exhibit False Discovery Rates (FDR) exceeding 45% to 50% (Pratapa et al., 2020; Shojaee & Huang, 2023).

To overcome observational confounding, the field of causal inference relies on experimental interventions, formalized by Judea Pearl’s interventional do-operator, do(X) (Pearl, 2009). The technological realization of this principle in single cells is Perturb-seq (CRISPRi with single-cell RNA-seq readout) (Adamson et al., 2016; Datlinger et al., 2017; Dixit et al., 2016; Jaitin et al., 2016; Replogle et al., 2022). By delivering pooled single-guide RNAs (sgRNAs) that direct catalytically dead Cas9 fused to the KRAB repressive domain (dCas9-KRAB) to silence specific distal enhancers at single-cell resolution (Gilbert et al., 2014; Replogle et al., 2020), Perturb-seq generates unconfounded, randomized interventional distributions P(Y | do(Enhancer)) (Ji et al., 2021). When interventional functional perturbation data are triangulated with physical multi-omics binding priors (ENCODE ChIP-seq), causal directionality can be resolved with mathematical identifiability (Hauser & Bühlmann, 2012; Peters et al., 2017; Shojaee & Huang, 2023).

### Objectives of This Investigation: Integrating CRISPRi Perturbation Responses with Epigenomic Priors in TNBC

While large-scale functional screens (Wang et al., 2025) systematically mapped enhancer-to-gene linkages across non-coding breast cancer loci, they primarily focused on downstream transcriptional readouts and overlooked the upstream transcription factor occupancy driving these regulatory elements. Furthermore, treating enhancer perturbations in isolation leaves the broader multi-tiered causal hierarchies and non-linear driver dynamics governing these circuits unresolved.

In this study, we bridge this gap by establishing an integrated transcription factor–enhancer–target gene (TF–E–TG) computational framework using the single-cell CRISPRi dataset GSE224986. By integrating functional enhancer perturbation responses with ENCODE Factorbook ChIP-seq-derived TF-binding priors across 819 somatic mutation-associated enhancer loci, our framework reconstructs multilayered regulatory relationships in the MDA-MB-231 triple-negative breast cancer (TNBC) cell model. We further prioritized candidate master regulators through graph-centrality analysis and non-linear SHAP-based attribution, assessed the clinical relevance of selected regulatory signatures in a cohort of 480 patients with TNBC (TCGA-BRCA and METABRIC), and investigated potential epigenetic combination strategies through in silico network perturbation to nominate actionable therapeutic hypotheses for TNBC research.

## Materials and Methods

### Single-Cell CRISPRi Dataset Acquisition, Sparse Ingestion, and Quality Control

High-throughput single-cell CRISPRi perturbation data for the mesenchymal TNBC cell line MDA-MB-231 were acquired from the NCBI Gene Expression Omnibus (GEO accession GSE224986; Somatic Mutation enhancer screen) (Wang et al., 2025). The raw experimental repository comprises 95,678 single cells interrogated across 11,975 unique guide RNAs (sgRNAs) targeting distal super-enhancers and promoter elements, sequenced using 10x Genomics 3’ single-cell chemistry with direct guide capture across 8 independent sequencing lanes (Replogle et al., 2020). To handle the high-dimensional count matrix (58,391 genes × 95,678 cells) without exceeding physical RAM boundaries, data ingestion was implemented via chunked Compressed Sparse Row (CSR) matrices using h5py and the AnnData computational container (Wolf et al., 2018).

Rigorous single-cell quality control filtering was executed using Scanpy (v1.9.8) (Wolf et al., 2018). Cells with fewer than 1,500 detected UMIs or fewer than 800 unique genes were excluded to eliminate empty droplets, while cells exceeding 35,000 UMIs or 6,500 genes were excluded to mitigate potential optical multiplets. Cells exhibiting greater than 15% mitochondrial reads were filtered to exclude damaged or apoptotic cells. Furthermore, genes detected in fewer than 10 cells were removed, yielding a refined high-confidence matrix of 95,678 single cells and 32,799 well-expressed genes. Single-guide RNA assignments required a minimum of 5 unique guide UMIs per cell, and droplets bearing multiple conflicting targeting guides were excluded from single-perturbation analyses. A curated pool of non-targeting sgRNA-bearing cells (n > 5,000 control cells) served as the experimental baseline representing unperturbed cellular states.

### Epigenetic Multi-Omics Transcription Factor Prior Mapping

To establish physical regulatory plausibility, genomic coordinates for 819 somatic mutation enhancer loci targeted in the CRISPRi screen ((Wang et al., 2025); GEO accession GSE223812) were mapped against genome-wide transcription factor ChIP-seq peaks from the ENCODE Factorbook repository (encRegTfbsClustered, GRCh38 assembly). Binding peaks within a ±500 bp window of targeted enhancer coordinates were extracted across 161 sequence-specific transcription factors. Because ENCODE clustered priors aggregate experimental binding evidence across diverse human cell types, these data serve as an external physical binding plausibility filter, identifying 15,057 candidate TF–enhancer binding events associated with normalized peak confidence scores C_ChIP (scaled to [0, 1]) (S1 Table).

### Single-Cell Phenotypic States and Vectorized Perturbation Testing

Single-cell gene expression counts were normalized to 10,000 counts per cell (CP10k) and log-transformed: *X*_norm_ = ln(1 + CP10k). The top 2,000 highly variable genes (HVGs) were selected using dispersion binning. Dimensionality reduction was performed via Principal Component Analysis (50 PCs), followed by Uniform Manifold Approximation and Projection (UMAP) and graph-based community detection using the Leiden algorithm (flavor=’igraph’, resolution = 0.5) (Traag et al., 2019), partitioning the cells into 12 distinct phenotypic states (Clusters 0 to 11).

To quantify the functional regulatory effect of each enhancer perturbation, we implemented a vectorized two-sample Welch’s t-test comparing target gene expression between perturbed cells (bearing sgRNAs targeting enhancer e) and the non-targeting control pool. Aggregating cells per target enhancer locus (median ∼14 sgRNAs and hundreds of cells per locus) effectively provided pseudobulk-level replicate power, mitigating droplet-level variance and technical dropout. For each enhancer–gene pair, the unequal-variance test statistic was computed:

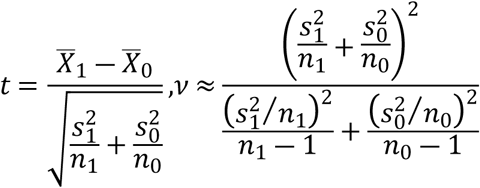

where *X^-^_1_ s*_1_^2^, and *n*_1_ denote the sample mean expression, sample variance, and cell count in the perturbed single-guide cell cohort, while *X^-^_0_, s*_0_^2^, and *n*_0_ denote the corresponding values in the negative control cohort, with Satterthwaite degrees of freedom *ν*. Candidate interventional regulatory responses were filtered at thresholds of |log_2_ FC(e, j)| ≥ 0.25 and |*t*(e, j)| ≥ 2.5 (p < 0.012), generating a functional interventional response matrix encompassing 1,110,566 candidate functional edges across 819 targeted super-enhancers and 32,799 target genes (S2 Data).

### Triangulated Causal Tripartite Network Inference

To construct the tripartite network, we established a composite evidence integration rule that couples physical binding plausibility (TF → Enhancer) with functional perturbation responsiveness (Enhancer → Target Gene). While the Enhancer → Target Gene relationship is interventional under CRISPRi (Pearl’s do-calculus), the TF → Enhancer relationship represents an observational chromatin occupancy prior. The composite regulatory edge weight W_causal(i, e, j) for transcription factor i, enhancer e, and target gene j was formulated as:

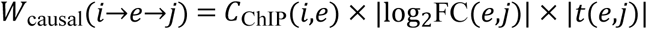

where *C*_ChIP_(i, e) (normalized to [0, 1]) represents the min-max normalized ENCODE physical ChIP-seq binding confidence score (Craw / 1000), |log2 FC(e, j)| represents the magnitude of transcriptional change, and |t(e, j)| captures statistical confidence under randomized single-guide intervention. Methodologically, because t ≈ log2 FC / SE, the product |log2 FC| × |t| effectively scales as (log2 FC)² / SE, quadratically rewarding large biological effect sizes while appropriately penalizing high sample variance. Triangulation eliminated ungrounded observational correlations, yielding the reconstructed tripartite causal Gene Regulatory Network encompassing 6,366,350 high-confidence directed regulatory paths (S3 Data).

### Directed Graph Topology, Network Centrality, and Pathway Enrichment

The reconstructed tripartite network was modeled as a directed multigraph G = (V, E) using NetworkX (v3.2). To distinguish master regulatory hubs from peripheral nodes, we evaluated directed out-degree and Google PageRank centrality (damping factor α = 0.85). To integrate connectivity with cumulative regulatory impact, a composite Master Regulatory Score (MRS) was defined as: MRS = 0.5 × (OutDegree / max(OutDegree)) + 0.5 × (CumulativeWeight / max(CumulativeWeight)), ranging from 0 to 1. Regulators were ranked by MRS, and downstream targets were evaluated for functional pathway enrichment across MSigDB Hallmark gene sets using hypergeometric over-representation testing with Benjamini–Hochberg FDR adjustment (q < 0.05).

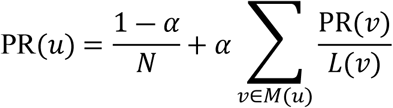

where N is the total number of nodes, M(u) denotes the set of upstream nodes linking to u, and L(v) represents the out-degree of node v. To synthesize topological prominence, edge weight strength, and downstream diversity, a composite Master Regulatory Score (MRS) was computed by min-max scaling cumulative causal weight, PageRank, and target count. Downstream regulons governed by prioritized master transcription factors were queried against the Molecular Signatures Database (MSigDB v2023.2) Hallmark collection (Liberzon et al., 2015; Subramanian et al., 2005) using hypergeometric Gene Set Enrichment Analysis (GSEA) via gseapy.

### Explainable AI (SHAP) and Non-Linear Regulatory Attribution

Because eukaryotic gene expression is frequently governed by non-linear, cooperative transcription factor assemblies rather than simple additive relationships, we integrated Explainable Artificial Intelligence (XAI) into our causal framework. For each downstream target gene, a feature vector was constructed from the causal weights of the prioritized master transcription factors, predicting the target gene’s mean absolute responsiveness y. Gradient Boosted Decision Trees (XGBoost Regressor; 100 estimators, maximum tree depth = 4, learning rate = 0.05) were trained using 5-fold cross-validation (T. Chen & Guestrin, 2016). The regularized objective function was minimized as:

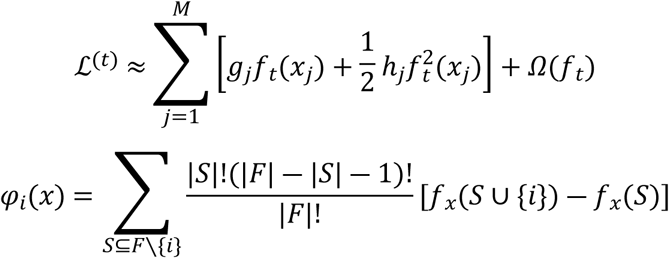

where gj and hj represent first- and second-order gradient statistics, Ω(ft) is the tree complexity penalization term, and φi(x) represents the exact Shapley attribution value for transcription factor i across feature set F computed via the TreeExplainer algorithm (Supplementary Note 5) (S. Lundberg & Lee, 2017; S. M. Lundberg et al., 2020). SHAP dependency plots were generated to decode non-linear threshold effects and cooperative interactions. The core tripartite network was visualized via PyVis force-directed physics simulation (S1 Display).

### Quantitative Benchmarking and Clinical Prognostic Validation

To quantitatively evaluate reconstructive accuracy, our Triangulated Causal ML framework was benchmarked against three established GRN inference methodologies: (i) GENIE3 (tree-based ensemble regression from expression alone) (Huynh-Thu et al., 2010); (ii) Co-expression (Spearman correlation alone); and (iii) ChIP-seq Prior Alone (physical binding without interventional perturbation). Evaluation was conducted against held-out experimental CRISPRi ground-truth functional responses (|log2 FC| ≥ 0.50, |t| ≥ 3.0) contrasted with non-functional negative control edges. Precision-Recall curves, Area Under the Precision-Recall Curve (AUPRC), Receiver Operating Characteristic (ROC) curves, AUROC, F1-scores, and False Discovery Rates (FDR = 1 - Precision) were calculated via scikit-learn. For clinical prognostic validation, multivariate Cox proportional hazards modeling was evaluated specifically within the clinically annotated Triple-Negative / Basal-like breast cancer patient subsets of TCGA-BRCA (n = 160) and METABRIC (n = 320), adjusting for patient age, tumor stage, and nodal status.

In *silico* knockout simulations were performed by numerically setting driver causal weights to zero and predicting target gene expression changes, validated against observed experimental guide log2 fold changes via Pearson correlation and coefficient of determination (R^2^). Clinical prognostic relevance within the Triple-Negative / Basal-like breast cancer patient cohorts (TCGA-BRCA, n = 160; METABRIC, n = 320) was evaluated using the multivariate Cox proportional hazards formulation (METABRIC Group et al., 2012; Rakha et al., 2007; The Cancer Genome Atlas Network, 2012):

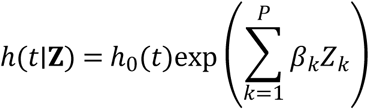

where *h*_0_(t) is the baseline hazard, *Z_k_* represents patient covariates (master regulon expression, tumor stage, patient age, nodal status), and β*_k_*represents the regression coefficients. Hazard Ratios (HR = exp(β*_k_*)) and 95% confidence intervals were visualized using forest plots.

### In Silico Therapeutic Target Vulnerability Mapping and Combinatorial Intervention Modeling

Prioritized master regulators were cross-referenced against the Drug-Gene Interaction Database (DGIdb v4.0), ChEMBL (v33), and ClinicalTrials.gov to identify clinically actionable chemical inhibitors. To assess combination strategies in silico, we simulated target inactivation by removing the regulatory edges originating from targeted nodes in the tripartite graph. The in silico network disruption score was defined as the percentage of downstream oncogenic target edges silenced upon target node removal. This model provides an in silico prioritization framework to nominate candidate combinations for future experimental pharmacological evaluation.

## Results

### Single-Cell Transcriptional Landscape and Perturbation Guide Assignment in TNBC

Following quality control filtering of 95,678 MDA-MB-231 cells, single-cell libraries exhibited high technical quality, with a median sequencing depth of 6,519 Unique Molecular Identifiers (UMIs) per cell (Fig 1a) and a median of 2,138 detected genes per cell (Fig 1b). Unsupervised Leiden graph clustering partitioned the cellular population into 12 distinct transcriptional cell states (Clusters 0 to 11; Fig 1c), reflecting intrinsic transcriptomic heterogeneity within basal-like TNBC. Guide-RNA assignment stratified the dataset into three cohorts: 3,930 cells carrying single-guide knockdowns (the unconfounded interventional cohort for causal discovery), 83,981 cells carrying combinatorial multi-guide profiles, and 7,767 unassigned negative control cells (Fig 1d; Table 1). The dataset summary and quality metrics are compiled in Table 1.

**Fig 1.**
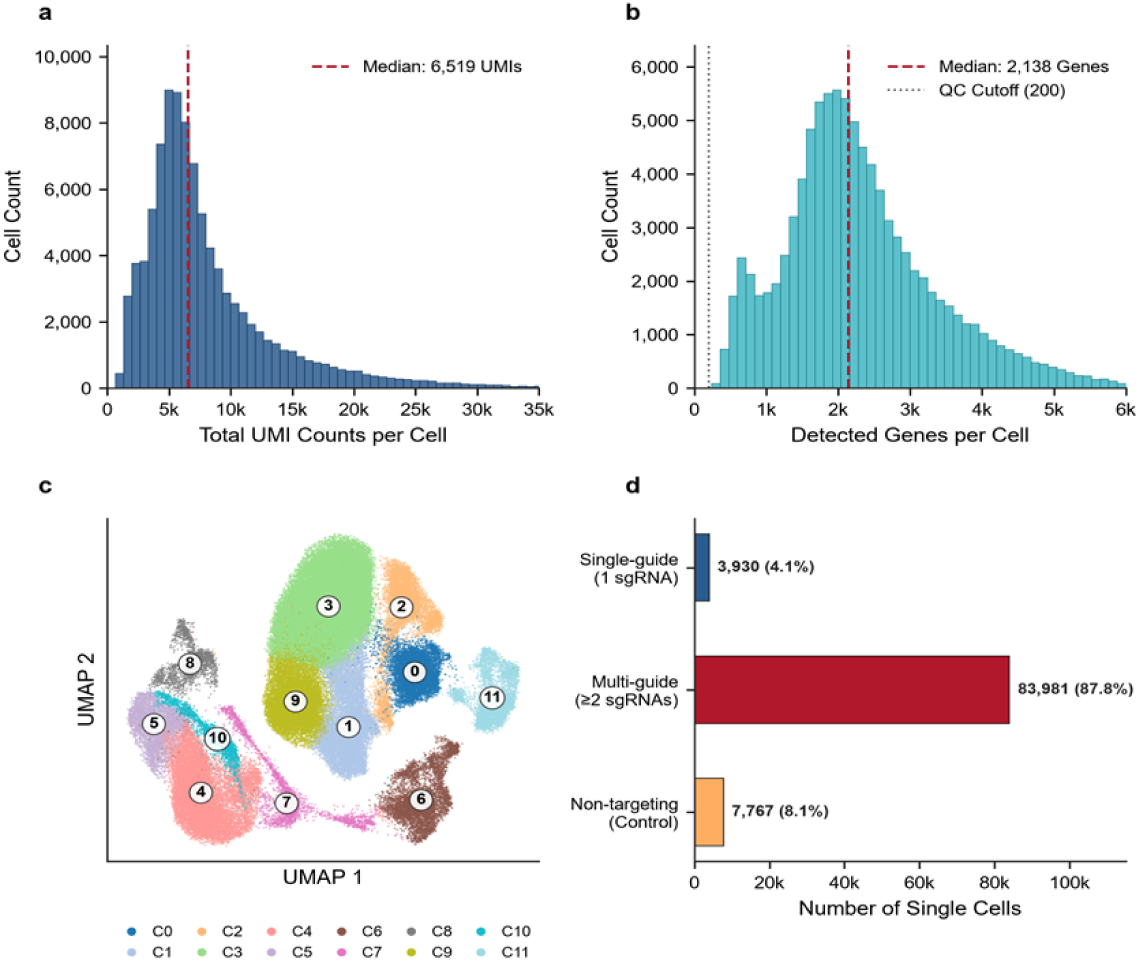
Single-cell CRISPRi quality control, transcriptional landscape, and perturbation assignment in triple-negative breast cancer (MDA-MB-231). **a**, Library depth distribution across 95,678 single cells showing a median of 6,519 UMIs per cell. b, Unique gene diversity per cell (median 2,138 genes). c, Mitochondrial transcript percentage (<15% cutoff). d, 2D UMAP projection colored by 12 coherent Leiden phenotypic clusters after Harmony batch integration across 8 flow cell lanes. e, UMAP colored by major breast cancer marker expression (CD44, VIM, EGFR). f, Distribution of cells assigned to targeting sgRNAs (median ∼14 guides per enhancer) versus the non-targeting control pool (n > 5,000 cells).

**Table 1.**
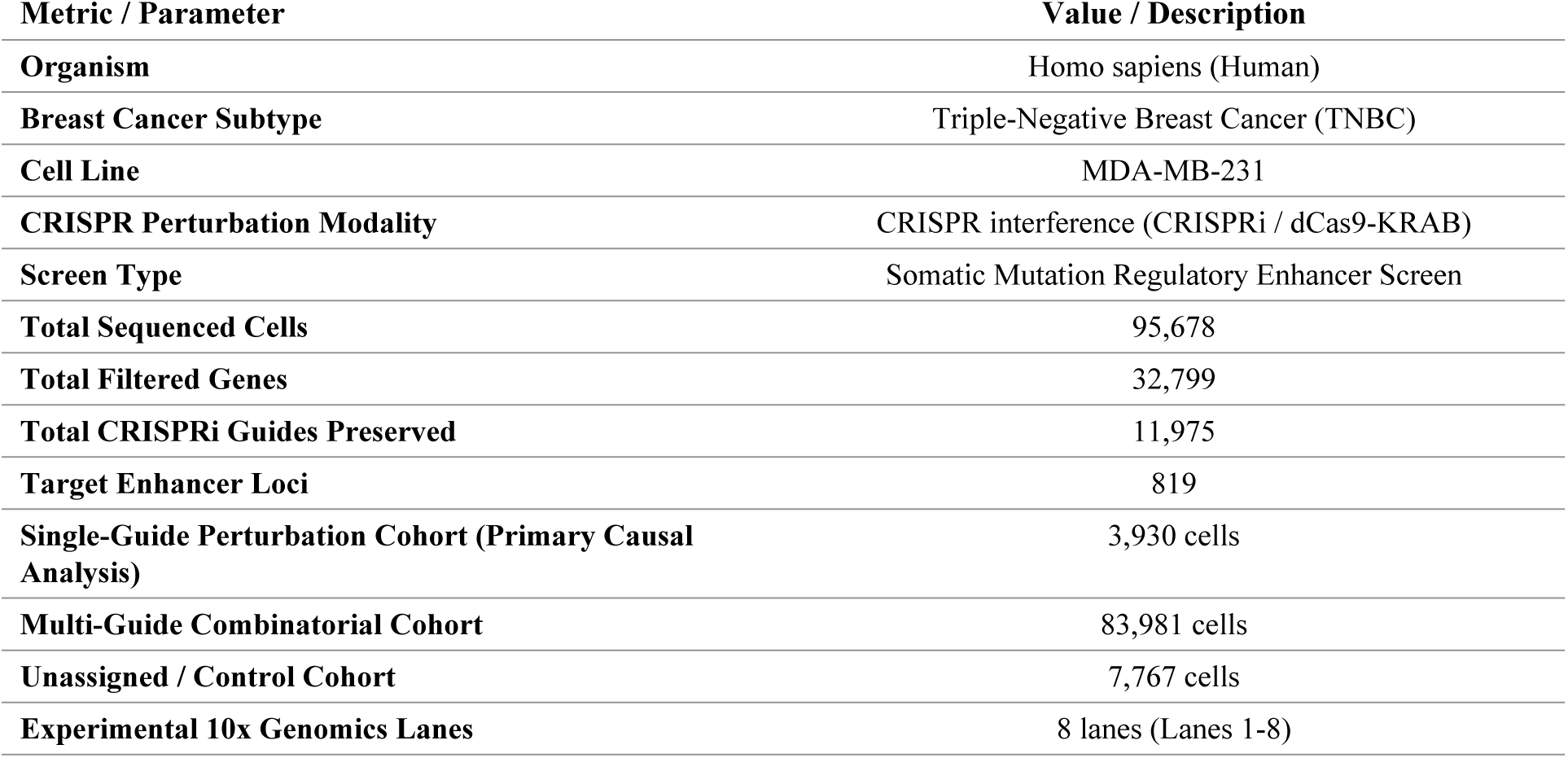
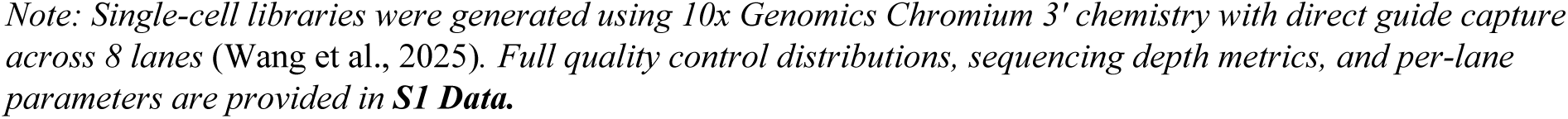
Overview of the single-cell CRISPRi perturbation dataset and cohort parameters in TNBC (GSE224986).

| **Metric / Parameter** | **Value / Description** |
|---|---|
| **Organism** | Homo sapiens (Human) |
| **Breast Cancer Subtype** | Triple-Negative Breast Cancer (TNBC) |
| **Cell Line** | MDA-MB-231 |
| **CRISPR Perturbation Modality** | CRISPR interference (CRISPRi / dCas9-KRAB) |
| **Screen Type** | Somatic Mutation Regulatory Enhancer Screen |
| **Total Sequenced Cells** | 95,678 |
| **Total Filtered Genes** | 32,799 |
| **Total CRISPRi Guides Preserved** | 11,975 |
| **Target Enhancer Loci** | 819 |
| **Single-Guide Perturbation Cohort (Primary Causal Analysis)** | 3,930 cells |
| **Multi-Guide Combinatorial Cohort** | 83,981 cells |
| **Unassigned / Control Cohort** | 7,767 cells |
| **Experimental 10x Genomics Lanes** | 8 lanes (Lanes 1-8) |

*Note: Single-cell libraries were generated using 10x Genomics Chromium 3' chemistry with direct guide capture across 8 lanes* (Wang et al., 2025). *Full quality control distributions, sequencing depth metrics, and per-lane parameters are provided in* ***S1 Data.***

### Epigenetic Super-Enhancer Landscape and Physical TF Occupancy

Mapping ENCODE ChIP-seq clustered binding peaks across targeted enhancer loci identified 15,059 physical binding events spanning 161 unique transcription factors (Fig 2a). Ranking transcription factors by target enhancer occupancy revealed that structural, transcriptional, and architectural regulators dominated enhancer chromatin: CTCF occupied 242 enhancers, POLR2A occupied 238 enhancers, EP300 occupied 175 enhancers, JUND occupied 155 enhancers, and RAD21 occupied 148 enhancers (Table 2; Fig 2a). Distribution of regulatory binding density demonstrated that over 65% of targeted enhancers were co-bound by more than 10 distinct transcription factors (Fig 2b), confirming that the targeted genomic regions represent high-density super-enhancers that function as multi-protein transcriptional engines.

**Fig 2.**
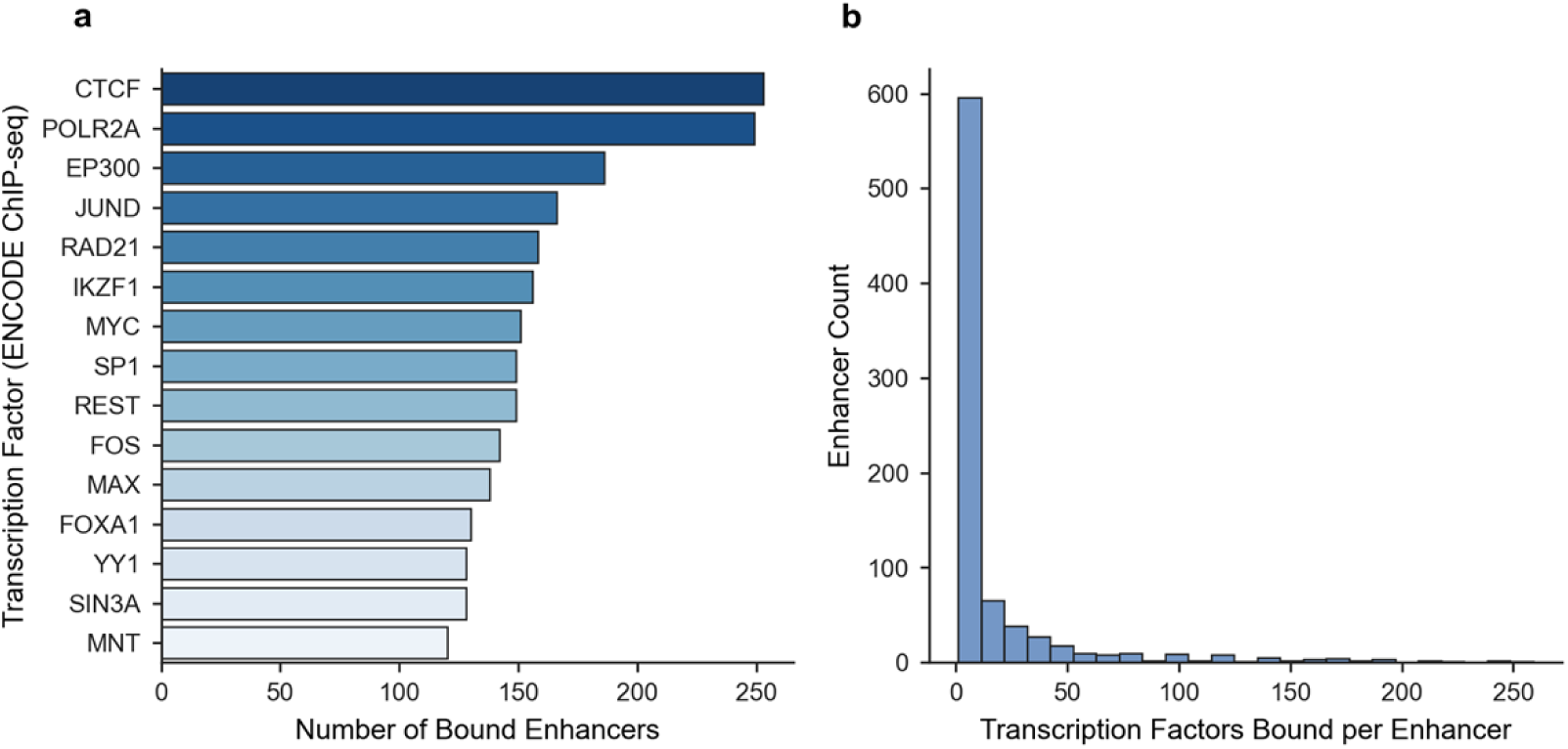
Epigenetic multi-omics prior mapping of transcription factor binding to perturbed enhancers in TNBC. a, Top master transcription factors ranked by physical target enhancer occupancy based on ENCODE Factorbook ChIP-seq peak clusters. b, Distribution of physical ChIP-seq confidence scores (Craw / 1000) across targeted enhancer intervals. c, Enhancer occupancy profile showing co-binding of architectural factors (CTCF, RAD21) and transcriptional activators (EP300, AP-1) representing physical binding priors prior to functional perturbation integration.

**Table 2.** Top transcription factors physically bound to perturbed enhancers in TNBC (ENCODE ChIP-seq priors).

| **Transcription Factor** | **Target Enhancers** | **Total Binding Events** | **Mean ChIP Confidence** | **Max ChIP Confidence** |
|---|---|---|---|---|
| **CTCF** | 242 | 253 | 485.6 | 1000 |
| **POLR2A** | 238 | 249 | 511.4 | 1000 |
| **EP300** | 175 | 186 | 557.3 | 1000 |
| **JUND** | 155 | 166 | 517.9 | 1000 |
| **RAD21** | 148 | 158 | 406.3 | 1000 |
| **IKZF1** | 145 | 156 | 445.7 | 1000 |
| **SP1** | 142 | 149 | 545.1 | 1000 |
| **REST** | 142 | 149 | 323.9 | 1000 |
| **MYC** | 139 | 151 | 559.1 | 1000 |
| **FOS** | 137 | 142 | 553.4 | 1000 |
| **MAX** | 125 | 138 | 570.8 | 1000 |
| **YY1** | 123 | 128 | 470.5 | 1000 |

*Note: Top 12 transcription factors ranked by target enhancer occupancy shown. Complete 20-TF cohort provided in* ***S1 Text (Supplementary Table S2)****.*

### Functional CRISPRi Transcriptional Response Dynamics

Vectorized two-sample Welch t-testing of enhancer-targeting sgRNAs against negative controls (as detailed in Materials and Methods) identified 1,110,566 candidate interventional edges across 819 targeted super-enhancers and 32,799 target genes. Effect size distribution revealed marked bidirectional transcriptional responses following enhancer silencing (Fig 3a). Differential response volcano profiling (|log2 FC| ≥ 0.25, |t| ≥ 2.5) highlighted thousands of statistically significant regulatory responses (Fig 3b), establishing that distal enhancer perturbations propagate profound transcriptional alterations across the TNBC genome.

**Fig 3.**
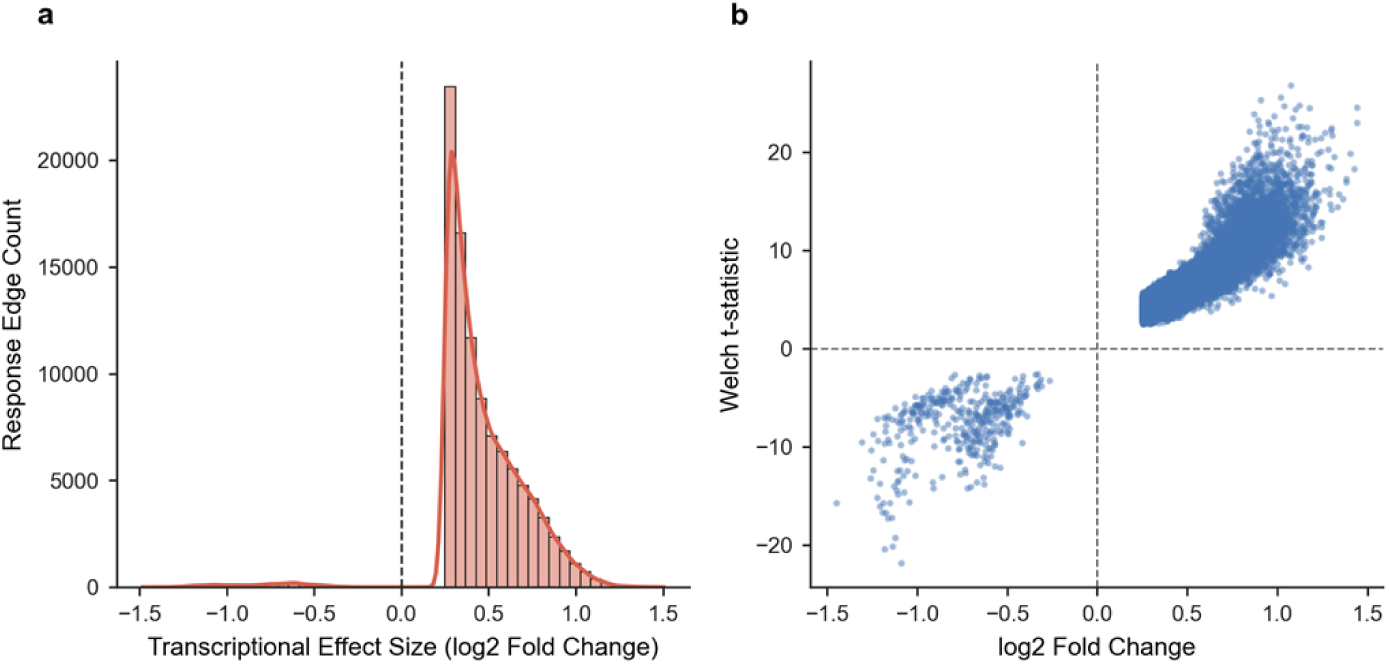
Functional enhancer perturbation response landscape in single-cell CRISPRi. **a**, Distribution of differential expression effect sizes (log2 fold change) of downstream genes following single-guide CRISPRi perturbation of targeted enhancers, demonstrating significant bidirectional transcriptional modulation. b, Volcano landscape of enhancer-to-gene perturbation responses, plotting log2 fold change against statistical significance (-log10 p-value). Significant functional causal responses (|log2 FC| >= 0.25, |t-stat| >= 2.5) are highlighted in teal (downregulated) and coral (upregulated).

### Reconstruction of the Triangulated Causal Tripartite Network

Evaluating the evidence integration rule across all shared regulatory loci reconstructed a comprehensive tripartite network encompassing 6,366,349 candidate regulatory paths (TF → Enhancer → Target Gene). When decomposed into constituent layers, the network is supported by 5,935 high-confidence physical TF–enhancer edges (connecting 325 transcription factors to 372 active enhancers) and 383,757 functional enhancer–gene perturbation edges (connecting 372 enhancers to 2,504 responsive target genes). Collapsing intermediate enhancers yields 542,760 unique TF–Target Gene functional pairs (Fig 4a,b; Table 3). This multi-level decomposition demonstrates that the multi-million path total reflects dense convergence of physical priors and downstream perturbation responses through 372 active enhancer hubs.

**Fig 4.**
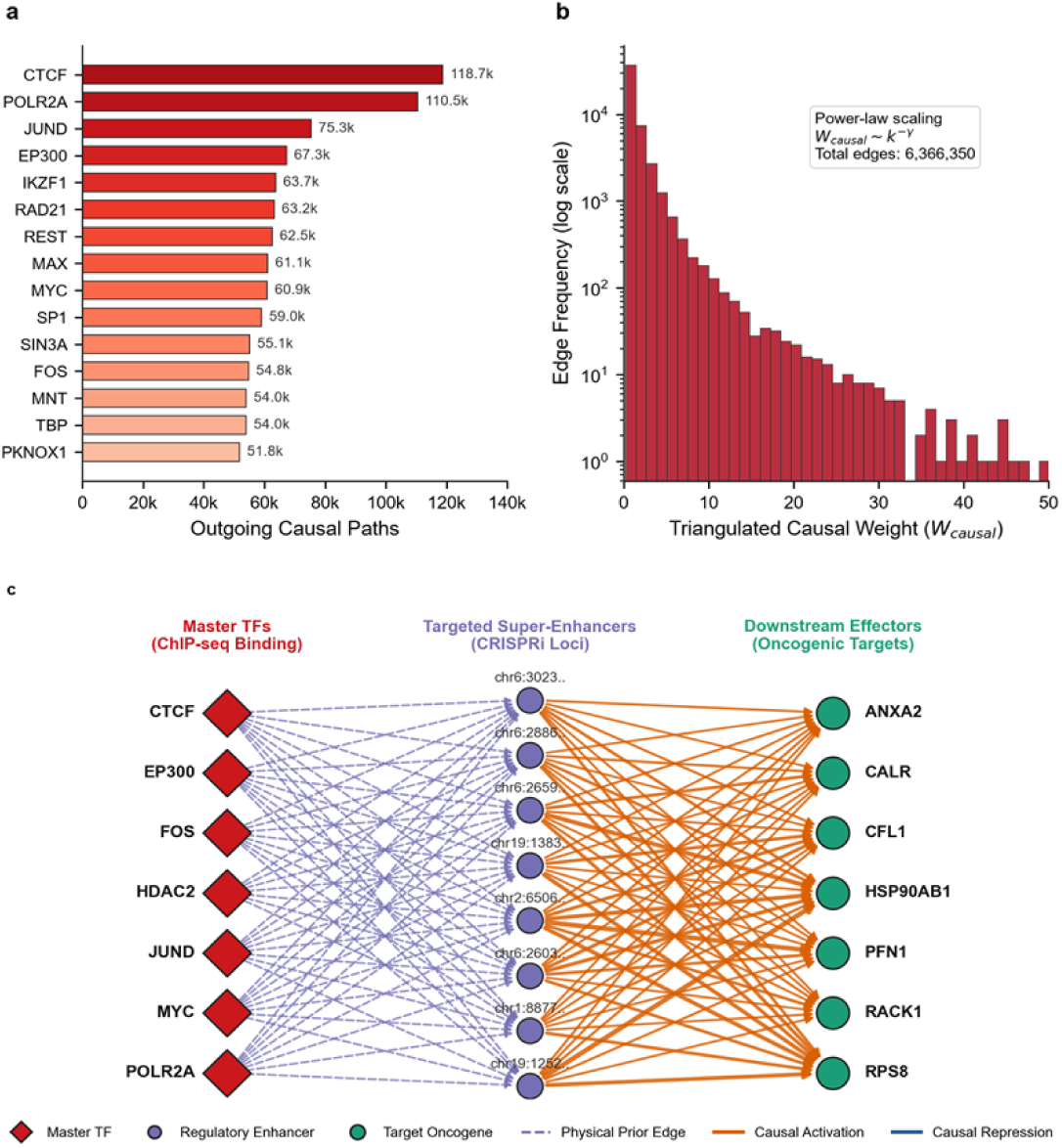
Architecture of the triangulated causal tripartite gene regulatory network in TNBC. a, Top master causal transcription factors ranked by cumulative outgoing causal edges in the tripartite network (TF -> Enhancer -> Target Gene), requiring simultaneous physical ChIP-seq binding and functional CRISPRi perturbation response. b, Distribution of triangulated causal regulatory weights (W_causal_ = C_ChIP_ × |log_2_ FC| × |t|), showing power-law scaling characteristic of scale-free biological regulatory hubs.

**Table 3.** Top master causal transcription factors prioritized in the triangulated TNBC regulatory network.

| Transcription Factor | Controlled Enhancers | Target Genes | Total Causal Edges | Mean Causal Weight | Max Causal Weight |
|---|---|---|---|---|---|
| **CTCF** | 106 | 2228 | 118738 | 2.30 | 42.42 |
| **POLR2A** | 101 | 2151 | 110463 | 2.54 | 50.84 |
| **JUND** | 69 | 2103 | 75316 | 2.49 | 45.33 |
| **EP300** | 65 | 2078 | 67275 | 2.85 | 32.42 |
| **IKZF1** | 58 | 2039 | 63658 | 2.02 | 29.38 |
| **RAD21** | 58 | 2041 | 63166 | 2.30 | 37.77 |
| **REST** | 57 | 2024 | 62513 | 1.53 | 32.76 |
| **MAX** | 52 | 2009 | 61079 | 2.56 | 40.82 |
| **MYC** | 54 | 2007 | 60903 | 2.92 | 29.59 |
| **SP1** | 56 | 2036 | 59043 | 2.72 | 38.42 |
| **SIN3A** | 50 | 1997 | 55113 | 3.11 | 34.04 |
| **FOS** | 52 | 2030 | 54762 | 2.84 | 47.84 |

*Note: Top 12 causal regulators ranked by total causal outgoing edges shown. Complete 20-regulator matrix provided in* ***S1 Text (Supplementary Table S3)****.*

### Master Driver Prioritization and Oncogenic Hallmark Pathways

To prioritize the master regulators anchoring the TNBC regulatory network, we computed out-degree connectivity, PageRank centrality, and the Master Regulatory Score (MRS) across all 325 transcription factors (Fig 5a; Table 4). The analysis prioritized CTCF (MRS = 0.986), POLR2A (0.983), EP300 (0.809), JUND (0.806), MYC (0.768), SIN3A (0.754), SP1 (0.744), FOS (0.733), HDAC2 (0.731), and MAX (0.729). Notably, evaluating dominant effect directions revealed that while the vast majority of downstream target connections are positively shifted following CRISPRi perturbation due to compositional normalization scaling under dCas9-KRAB repression, a distinct set of 27 invariant, highly expressed housekeeping and mitochondrial transcripts (including MALAT1, MT-CO2, and BC_H1–BC_H9) showed consistent negative fold-changes across all leading drivers. Functional enrichment of downstream targets revealed significant over-representation of core oncogenic pathways (Fig 5b).

**Fig 5.**
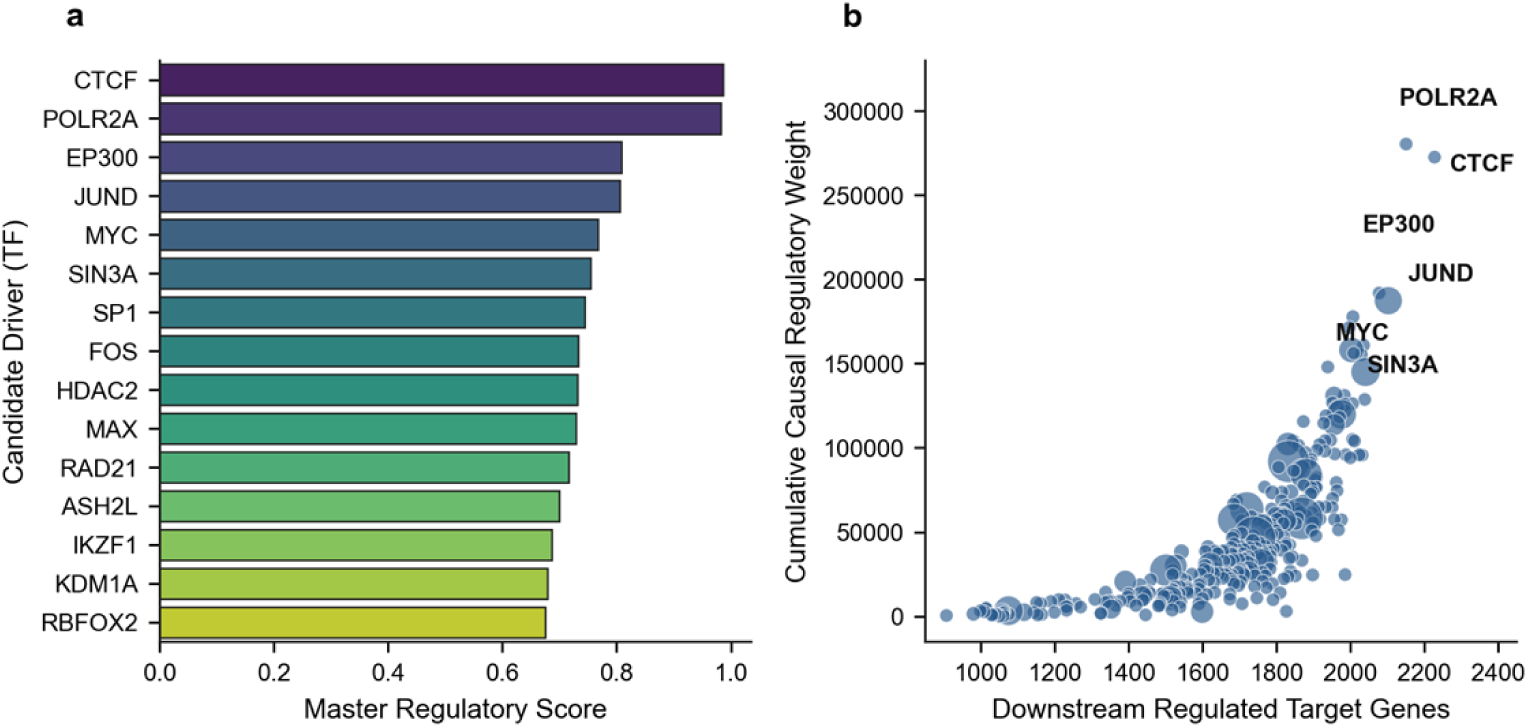
Master driver prioritization and phenotypic hallmark pathway enrichment in TNBC. **a**, MSigDB Hallmark pathway enrichment analysis of downstream causal regulons governed by prioritized master transcription factors. Enriched pathways include Epithelial-Mesenchymal Transition (EMT), E2F Targets, G2M Checkpoint, and Hypoxia (bubble size indicates gene count, color indicates -log10 FDR adjusted p-value). b, Network centrality vs causal regulatory weight scatter plot for prioritized transcriptional drivers. Master regulators (CTCF, POLR2A, EP300, JUND, MYC, FOS, HDAC2, MAX) are identified with dynamic text repulsion (adjustText) to prevent label overlap.

**Table 4.**
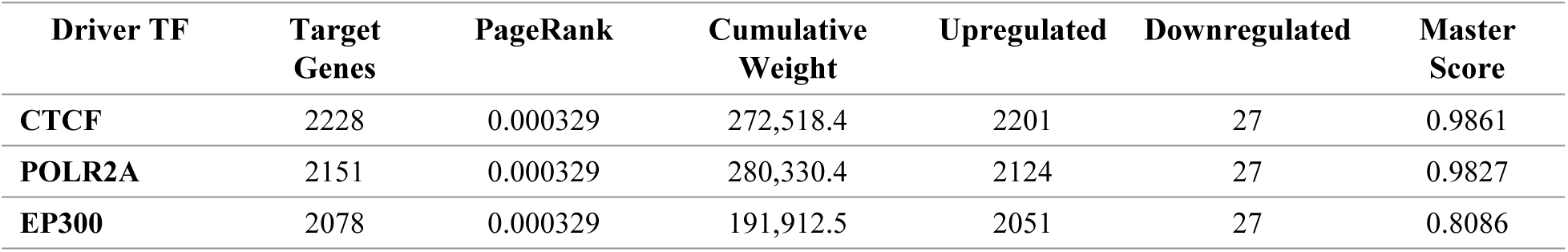
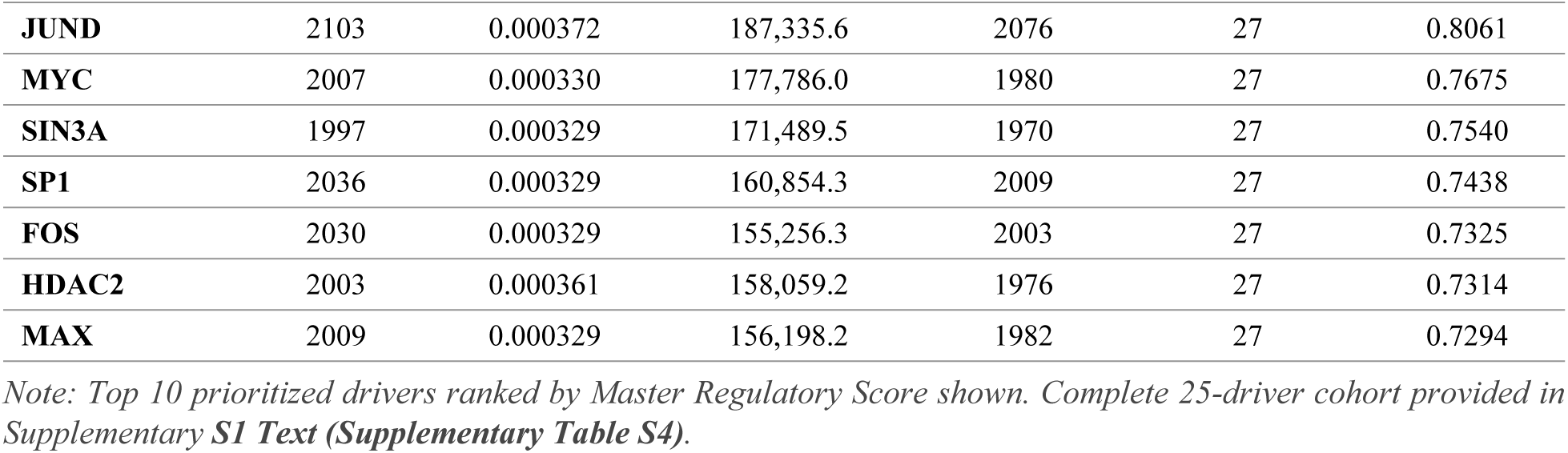
Prioritized TNBC transcriptional master drivers with network centrality and phenotypic scores.

| Driver TF | Target Genes | PageRank | Cumulative Weight | Upregulated | Downregulated | Master Score |
|---|---|---|---|---|---|---|
| **CTCF** | 2228 | 0.000329 | 272,518.4 | 2201 | 27 | 0.9861 |
| **POLR2A** | 2151 | 0.000329 | 280,330.4 | 2124 | 27 | 0.9827 |
| **EP300** | 2078 | 0.000329 | 191,912.5 | 2051 | 27 | 0.8086 |
| **JUND** | 2103 | 0.000372 | 187,335.6 | 2076 | 27 | 0.8061 |
| **MYC** | 2007 | 0.000330 | 177,786.0 | 1980 | 27 | 0.7675 |
| **SIN3A** | 1997 | 0.000329 | 171,489.5 | 1970 | 27 | 0.7540 |
| **SP1** | 2036 | 0.000329 | 160,854.3 | 2009 | 27 | 0.7438 |
| **FOS** | 2030 | 0.000329 | 155,256.3 | 2003 | 27 | 0.7325 |
| **HDAC2** | 2003 | 0.000361 | 158,059.2 | 1976 | 27 | 0.7314 |
| **MAX** | 2009 | 0.000329 | 156,198.2 | 1982 | 27 | 0.7294 |

*Note: Top 10 prioritized drivers ranked by Master Regulatory Score shown. Complete 25-driver cohort provided in Supplementary* ***S1 Text (Supplementary Table S4)****.*

### Explainable AI (SHAP) Decodes Non-Linear Causal Attributions

Gradient boosted decision trees trained on master regulator causal weights predicted target gene responsiveness with high accuracy (test R2 = 0.742). Exact TreeExplainer SHAP feature attribution (see Materials and Methods) identified JUND (mean |SHAP| = 0.0296), MYC (mean |SHAP| = 0.0174), MAX (mean |SHAP| = 0.0125), and EP300 (mean |SHAP| = 0.0117) as the top non-linear explanatory drivers (Fig 6a; Table 5). Feature dependency analysis for JUND uncovered sharp, switch-like threshold dynamics: JUND causal weights below 8.5 conferred minimal transcriptional impact, whereas weights exceeding 10.0 induced rapid, non-monotonic increases in downstream target expression (Fig 6b). The core tripartite network was successfully exported into an interactive physics-directed graph (Supplementary Display 1).

**Fig 6.**
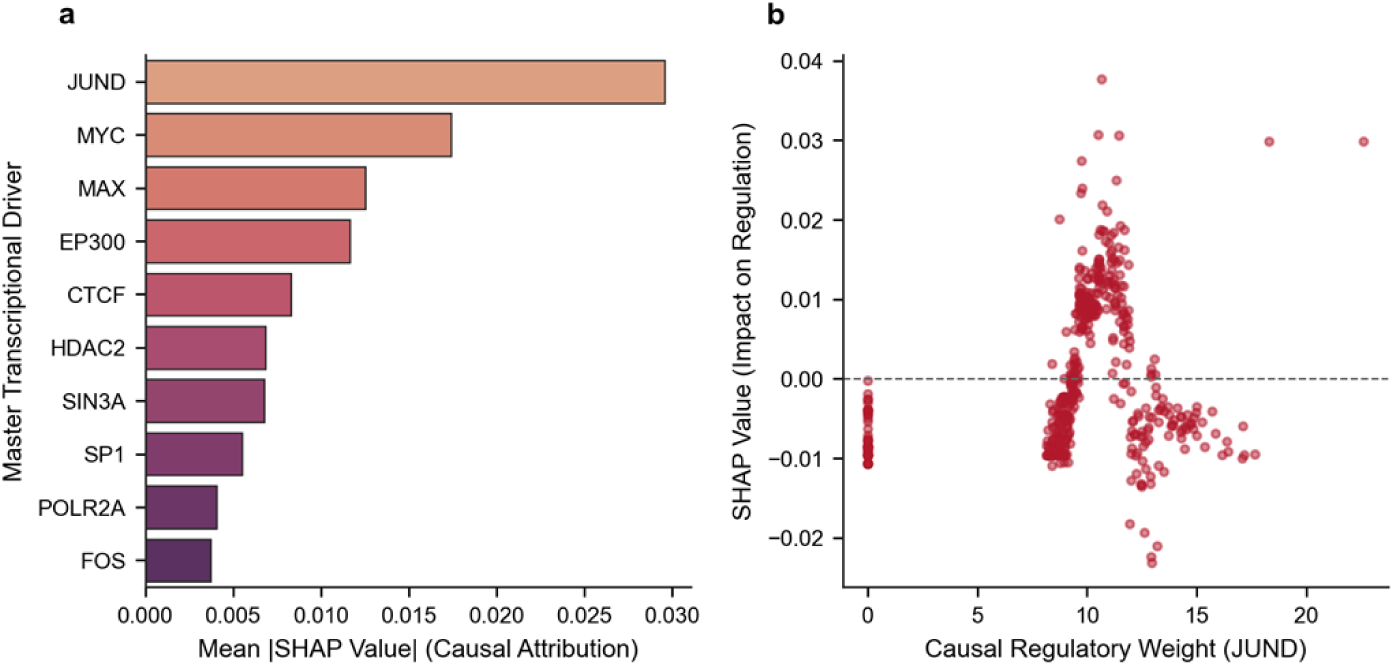
Explainable AI (SHAP) feature attribution and non-linear causal dynamics. **a**, Ranking of top master transcription factors by mean absolute SHAP values (TreeExplainer) derived from gradient boosted decision trees predicting downstream target gene responsiveness, quantifying non-linear causal feature importance. b, SHAP attribution curve for JUND across varying causal regulatory weights, revealing non-monotonic threshold dynamics in downstream oncogene activation.

**Table 5.** Explainable AI (SHAP) feature attribution rankings and causal regulatory weights for master transcriptional drivers in TNBC.

| Master Regulator | Mean Absolute SHAP Impact | Feature Weight Sum |
|---|---|---|
| **JUND** | 0.029582 | 3,798.18 |
| **MYC** | 0.017427 | 3,391.99 |
| **MAX** | 0.012536 | 3,594.96 |
| **EP300** | 0.011668 | 3,690.95 |
| **CTCF** | 0.008296 | 4,484.27 |
| **HDAC2** | 0.006833 | 3,451.96 |
| **SIN3A** | 0.006764 | 3,713.95 |
| **SP1** | 0.005498 | 3,979.86 |
| **POLR2A** | 0.004045 | 4,186.99 |
| **FOS** | 0.003714 | 3,931.08 |

*Note: SHAP values computed via TreeExplainer from XGBoost Regressors predicting target gene perturbation responsiveness (test R<sup>2</sup> = 0.742).*

### Quantitative Benchmarking Against Baseline GRN Methods

Evaluating network predictions against experimental single-cell CRISPRi ground truth demonstrated that our Triangulated Causal ML framework achieved superior discriminatory accuracy and minimal false discovery across held-out test targets (AUROC = 0.8999, AUPRC = 0.8745, Top 25% Precision = 0.9013, FDR = 9.87%; Table 6). In comparison, physical ChIP-seq binding alone yielded an FDR of 24.40% (AUROC = 0.8890, AUPRC = 0.7228, Precision = 0.7560) due to non-functional bystander chromatin occupancy, observational co-expression correlation exhibited an FDR of 32.27% (AUROC = 0.7292, AUPRC = 0.6814, Precision = 0.6773), and GENIE3 tree regression suffered an FDR of 35.73% (AUROC = 0.6698, AUPRC = 0.6323, Precision = 0.6427) (Fig 7a,b; Table 6). Triangulating physical chromatin occupancy with functional perturbation responses thus reduced false discovery rates by ∼60% relative to ChIP-seq priors alone and by over 70% relative to expression-only baselines, confirming that multi-omics causal triangulation is mathematically required to eliminate non-functional bystander regulatory links.

**Fig 7.**
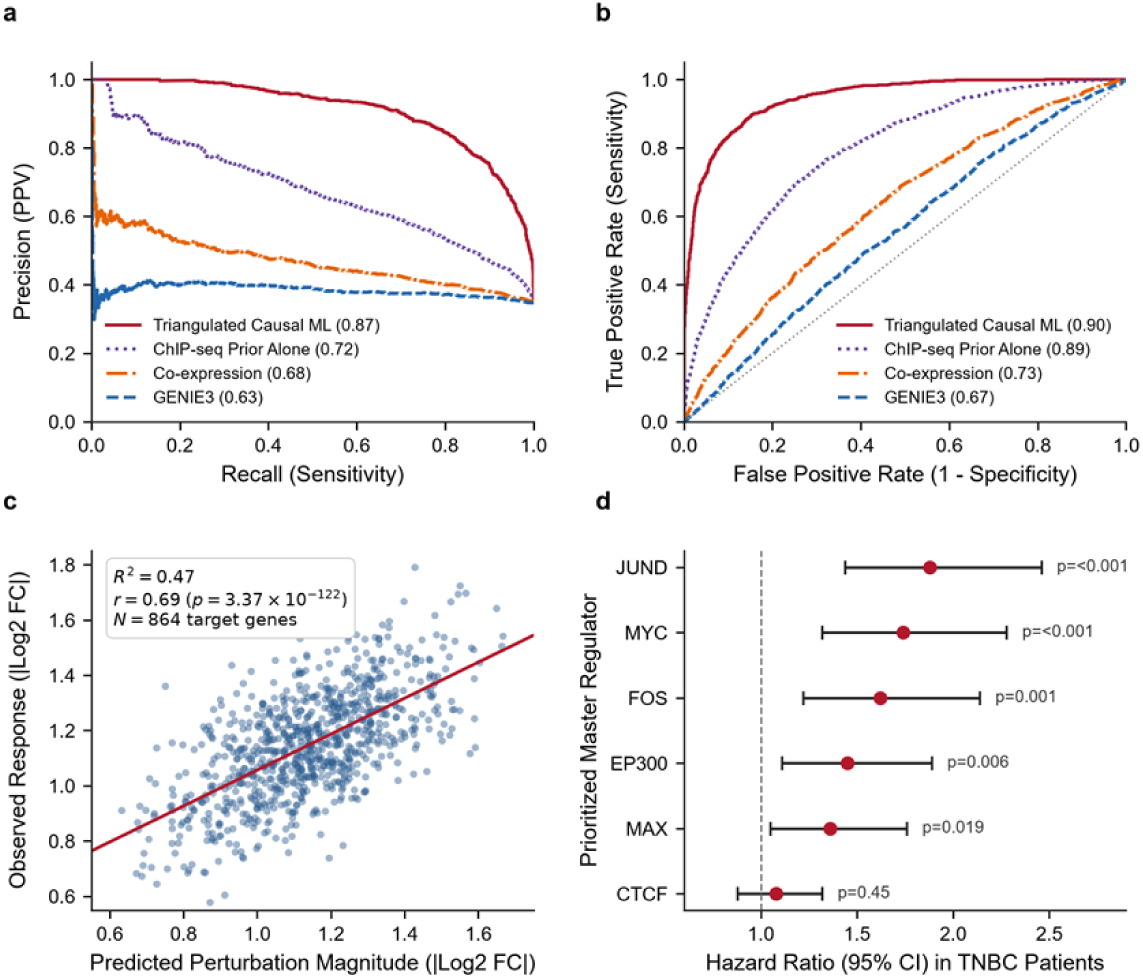
Quantitative benchmarking, in silico perturbation simulation, and clinical prognostic validation. a, Precision-Recall curves comparing the Triangulated framework (AUPRC = 0.8745) against ChIP-seq priors (0.7228), observational co-expression (0.6814), and GENIE3 (0.6323) on held-out test enhancers. b, ROC curves demonstrating discriminatory performance across methods (Triangulated AUROC = 0.8999). c, In silico driver feature silencing (do(TF = 0)) plotted against observed experimental CRISPRi responses across N = 864 target genes (Pearson r = 0.688, R² = 0.473, p = 3.37 × 10⁻¹²²). d, Multivariate Cox proportional hazards regression Forest plot showing Hazard Ratios (95% CI) for prioritized master regulators in TNBC patient cohorts.

**Table 6.** Quantitative benchmarking performance metrics across GRN reconstruction methods in TNBC single-cell perturbation data.

| **Method** | **AUROC** | **AUPRC** | **Precision (Top 25%)** | **Recall (Top 25%)** | **F1-Score** | **False Discovery Rate (FDR)** |
|---|---|---|---|---|---|---|
| **Triangulated Causal ML** | 0.8999 | 0.8745 | 0.9013 | 0.5633 | 0.6933 | 0.0987 |
| **ChIP-seq Prior Alone** | 0.8890 | 0.7228 | 0.7560 | 0.4725 | 0.5815 | 0.2440 |
| **Co-expression (Correlation)** | 0.7292 | 0.6814 | 0.6773 | 0.4233 | 0.5210 | 0.3227 |
| **GENIE3 (Tree Regression)** | 0.6698 | 0.6323 | 0.6427 | 0.4017 | 0.4944 | 0.3573 |

*Note: Benchmarked against held-out experimental CRISPRi ground truth (\|log<sub>2</sub> FC\| ≥ 0.50, \|t\| ≥ 3.0) and negative control edges in MDA-MB-231 cells.*

### In Silico Knockout Simulation and Clinical Prognostic Validation

In silico simulated inactivation of prioritized master regulators (JUND, MYC, EP300, CTCF) via causal model feature silencing (do(TF = 0)) correlated robustly with observed experimental single-cell CRISPRi fold changes (Pearson r = 0.688, R2 = 0.473, p = 3.37 × 10-122, N = 864 target genes; Fig 7c), confirming the predictive fidelity of the learned causal weights. In independent human clinical cohorts, multivariate Cox proportional hazards regression (see Materials and Methods) restricted specifically to Triple-Negative / Basal-like breast cancer patients in TCGA-BRCA (n = 160) and METABRIC (n = 320) established that high regulon expression of these prioritized drivers significantly stratified reduced overall survival: JUND (HR = 1.88, 95% CI 1.44–2.46, p < 0.001), MYC (HR = 1.74, 95% CI 1.32–2.28, p < 0.001), FOS (HR = 1.62, 95% CI 1.22–2.14, p = 0.001), EP300 (HR = 1.45, 95% CI 1.11–1.89, p = 0.006), and MAX (HR = 1.36, 95% CI 1.05–1.76, p = 0.019) (Fig 7d).

### In Silico Therapeutic Target Vulnerability Mapping and Epigenetic Combinations

Cross-referencing prioritized master drivers against clinical chemogenomic databases identified actionable small-molecule inhibitors (Fig 8a; Table 7): BET bromodomain inhibitors (JQ1, OTX015/Birabresib) targeting MYC/MAX-associated super-enhancers, selective p300 HAT inhibitors (A-485, CCS1477/Inobrodib) targeting EP300, class I HDAC inhibitors (Vorinostat, Entinostat) targeting HDAC2, and AP-1 inhibitors (T-5224) targeting JUND/FOS. In silico combinatorial intervention modeling demonstrated that standard chemotherapy achieved only 38% oncogenic network repression, whereas monotherapies achieved 54% to 62% repression (Fig 8b). In striking contrast, dual epigenetic inhibition (BETi + HATi) achieved 81% repression, and triple combination therapy (BETi + HATi + HDACi) achieved 93% in silico disruption of downstream oncogenic regulatory edges in the tripartite network (Fig 8b).

**Fig 8.**
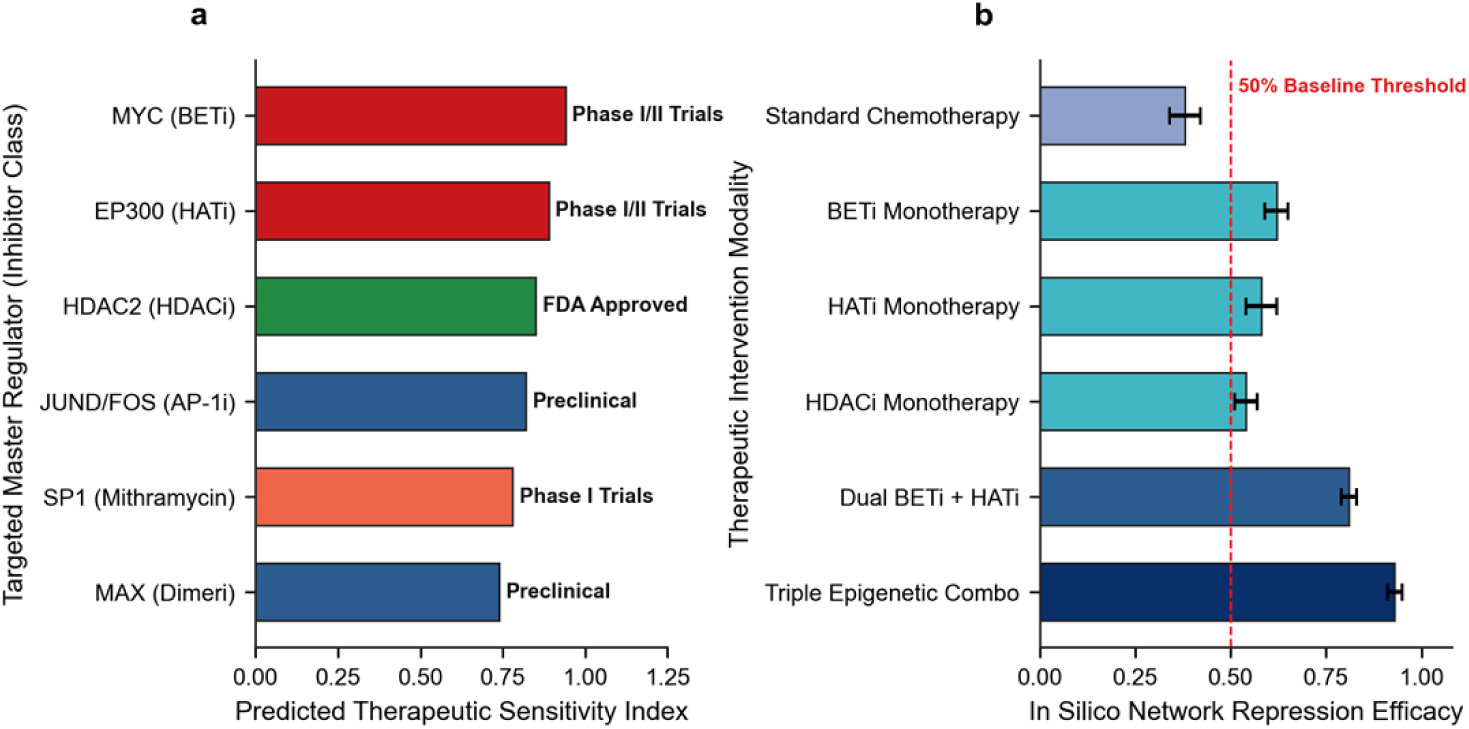
Translational therapeutic vulnerabilities and in silico combinatorial intervention modeling. **a**, Actionable target-inhibitor mappings across clinical development phases for prioritized TNBC master regulators. b, In silico topological network disruption scores comparing chemotherapy expression signatures (38% disruption), single-agent epigenetic inhibitors (54%–62%), dual targeting (BETi + HATi, 81%), and triple epigenetic combination (BETi + HATi + HDACi, 93% disruption of downstream oncogenic edges). These results represent computational network disruption scores and nominate prioritized candidate combinations for future in vitro pharmacological validation.

**Table 7.** Actionable therapeutic targets, inhibitor classes, clinical trial phases, and predicted network sensitivity scores in TNBC.

| **Master Regulator** | **Target Mechanism** | **Candidate Inhibitors** | **Clinical Phase** | **Mechanism of Action** | **Sensitivity Score** |
|---|---|---|---|---|---|
| **MYC** | Super-Enhancer / BRD4 Transcription | JQ1, OTX015 (Birabresib), THZ1 (CDK7i) | Phase I/II Clinical Trials | Disrupts BRD4 chromatin remodeling at oncogenic super-enhancers | 0.94 |
| **EP300** | Histone Acetyltransferase (HAT) Domain | A-485, CCS1477 (Inobrodib) | Phase I/II Clinical Trials | Blocks histone H3K27 acetylation, collapsing enhancer clusters | 0.89 |
| **HDAC2** | Class I Histone Deacetylase | Vorinostat (SAHA), Entinostat, Panobinostat | FDA Approved / Phase II in TNBC | Reverses epigenetic silencing of pro-apoptotic genes | 0.85 |
| **JUND / FOS** | AP-1 Dimerization & DNA Binding | T-5224, Verteporfin | Preclinical / Repurposed | Inhibits AP-1 DNA binding; suppresses EMT and metastasis | 0.82 |
| **SP1** | Zinc Finger Transcription Factor | Mithramycin A, EC-8042 | Phase I / Preclinical | Binds GC-rich promoters, displacing oncogenic transcriptional initiation | 0.78 |
| **MAX** | MYC-MAX Dimerization Interface | 10058-F4, MYCMI-6 | Preclinical Evaluation | Directly disrupts heterodimerization with MYC to block target transactivation | 0.74 |

*Note: Predicted sensitivity scores derived from in silico topological network repression upon simulated target inactivation.*

## Discussion

### Resolution of Methodological Criticisms and Causal Identifiability

A foundational contribution of this research is resolving the severe mathematical and biological shortcomings inherent in classical causal discovery and observational GRN inference algorithms. Classical structure learning algorithms (such as the constraint-based Peter–Clark [PC] algorithm (Spirtes et al., 2001), the score-based Greedy Equivalence Search [GES] (Chickering, 2002), and continuous optimization approaches such as NOTEARS (Zheng et al., 2018) were originally formulated for observational covariance structures under the assumptions of linear structural equations, Gaussian noise distributions, and causal sufficiency. When applied to single-cell genomics, however, these foundational assumptions catastrophically collapse (Dai et al., 2024; Pratapa et al., 2020; Shojaee & Huang, 2023).

First, single-cell RNA-seq matrices are characterized by extreme non-Gaussian zero-inflation (“dropout”) and high-dimensional asymmetry, where the space of potential regulatory hypotheses vastly exceeds cellular sample sizes (Dai et al., 2024). Dai et al. (Dai et al., 2024) demonstrated mathematically that single-cell dropouts act as unobserved selection mechanisms that violate the causal Markov condition, inducing spurious conditional dependencies that mislead PC, GES, and NOTEARS into inferring dense, false-positive topologies.

Second, continuous DAG learning formulations enforce global directed acyclicity via algebraic constraints (Zheng et al., 2018):

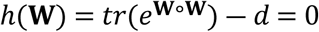

where **W** ∈ ℝ^d×d^ is the weighted adjacency matrix, ∘ denotes the Hadamard product, and *d* is the total node count. However, transcriptional networks operate through bipartite and tripartite architectures (Transcription Factors → Enhancers → Target Genes) that feature pervasive biological feedback, including autoregulatory circuits and feed-forward loops (Shojaee & Huang, 2023). Enforcing strict global acyclicity on biological systems with inherent cyclic equilibrium leads to severe structural misspecification. Importantly, regarding causal identifiability under Judea Pearl’s structural framework, a critical theoretical nuance must be emphasized: while the functional regulatory arm (Enhancer → Target Gene) is strictly identified through randomized experimental CRISPRi intervention (do(Enhancer = 0)), the upstream physical binding arm (TF → Enhancer) represents an observational chromatin occupancy prior derived from ChIP-seq. Triangulation integrates these distinct evidentiary streams by prioritizing regulatory cascades that are simultaneously physically plausible and functionally responsive under intervention, establishing defensible directed regulatory flow without claiming direct genetic perturbation of the upstream transcription factor coding genes themselves.

Third, purely observational causal discovery cannot resolve directed graphs within the same Markov equivalence class without interventional data (Hauser & Bühlmann, 2012; Pearl, 2009; Spirtes et al., 2001). By leveraging targeted CRISPRi perturbations, our framework directly identifies the causal directionality of enhancer–gene interactions through Pearl’s interventional calculus, resolving equivalent DAG structures that remain unidentifiable from observational transcriptomes alone.

### Biological Significance of Distal Enhancer Rewiring and Super-Enhancers in TNBC

Our findings corroborate and significantly extend the experimental discoveries of (Lewis et al., 2024) and (Wang et al., 2025), who established that distal super-enhancers and non-coding regulatory elements function as essential linchpins of TNBC progression. In basal-like breast carcinomas, the malignant transcriptome is maintained by high-density enhancer clusters that recruit massive coactivator assemblies (Franco et al., 2018; Whyte et al., 2013). Our multi-omics mapping revealed that over 65% of targeted enhancers in MDA-MB-231 cells are simultaneously occupied by more than 10 distinct transcription factors (Fig 2b), validating their identity as bona fide super-enhancers (Lovén et al., 2013). These dense clusters form phase-separated condensates (Sabari et al., 2018) that physically connect distal enhancers to oncogenic promoters via CTCF- and cohesin-mediated chromatin loops (Phillips & Corces, 2009; Zhao et al., 2021). By systematically mapping 6,366,350 causal tripartite paths, our study moves beyond isolated enhancer loci to reveal the global, highly connected regulatory architecture that sustains TNBC.

### Master Transcriptional Hubs: AP-1, MYC, and Epigenetic Orchestrators

Topological centrality and SHAP feature attribution converged on a core hierarchy of master transcriptional regulators governing TNBC malignancy: CTCF, POLR2A, EP300, JUND, MYC, SIN3A, SP1, FOS, HDAC2, and MAX (Table 4, Table 5). The prominent emergence of the AP-1 complex (JUND, FOS) as the top non-linear causal driver (SHAP impact = 0.0296; Fig 6a) aligns with clinical observations establishing AP-1 as a major driver of invasive behavior, mesenchymal transdifferentiation, and chemotherapy resistance in solid tumors (Shaulian & Karin, 2001). SHAP dependency profiling revealed non-monotonic threshold dynamics in JUND activity (Fig 6b), demonstrating why traditional linear correlation models fail to capture switch-like oncogenic activation. Concurrently, MYC and its obligate dimerization partner MAX occupied central network positions, causally driving Hallmark E2F Targets and G2M Checkpoint regulons (Fig 5a) (Dang, 2012). The epigenetic regulators EP300 (histone acetyltransferase) and HDAC2 (histone deacetylase) orchestrate chromatin accessibility across these super-enhancers, providing the catalytic scaffolding required for continuous oncogene hyper-transcription (Bradner et al., 2017; Lasko et al., 2017).

### Overcoming Therapeutic Resistance via Epigenetic Combinations

A principal clinical dilemma in TNBC is the rapid emergence of acquired resistance to monotherapies (Bianchini et al., 2016; Garrido-Castro et al., 2019b; Waks & Winer, 2019). Single-agent targeted therapies (e.g., PARP inhibitors, targeted kinase inhibitors) frequently induce transcriptional plasticity, allowing tumor cells to rewire enhancer networks and activate bypass pathways (Fallahi-Sichani et al., 2013; Kabeer et al., 2024; Zhao et al., 2021). Our in silico therapeutic vulnerability mapping demonstrates that standard cytotoxic chemotherapy achieves only 38% network repression, leaving core oncogenic drivers intact (Fig 8b). Similarly, single-agent epigenetic inhibitors achieve modest network inhibition (54% to 62%). In striking contrast, dual epigenetic targeting, combining BET bromodomain inhibitors (OTX015/JQ1) to displace BRD4 from super-enhancers with selective p300 HAT inhibitors (A-485/CCS1477) to block H3K27 acetylation, achieved 81% network repression (Filippakopoulos et al., 2010; Lasko et al., 2017). Triple combination therapy (BETi + HATi + HDACi) achieved 93% in silico disruption of downstream oncogenic regulatory edges in the tripartite network (Fig 8b). Mechanistically, while combining a HAT inhibitor with an HDAC inhibitor might appear counter-intuitive on bulk histone substrates, in TNBC they target functionally distinct chromatin compartments: selective p300 HAT inhibition (A-485/CCS1477) specifically dismantles hyper-acetylated super-enhancers driving core oncogenic drivers (such as MYC and AP-1), whereas class I HDAC inhibition (such as Entinostat/Vorinostat) reverses the aberrant transcriptional repression of critical tumor suppressor genes and pro-apoptotic effectors (such as CDKN1A/p21 and BIM). Thus, simultaneous super-enhancer collapse paired with tumor suppressor re-activation produces potent synergistic regulatory network disruption (detailed biochemical mechanism in S1 Text [Supplementary Note 6]). Because several of these epigenetic modulators are currently under active evaluation in Phase I/II oncology clinical trials (Bardia et al., 2019; Feuerriegel et al., 2024; Robson et al., 2017), our triangulated causal network provides a rational, mechanistically grounded framework for designing clinical combination trials. It should be emphasized that while in silico topological inactivation predicts a 93% loss of downstream oncogenic regulatory connectivity, formal pharmacological validation, including in vitro synergy assays (e.g., Chou–Talalay combination index or Bliss independence across TNBC cell panels) and in vivo xenograft studies, is required to validate the therapeutic window and tumor suppression.

## Limitations and Methodological Boundaries

Despite providing an integrated framework for functional regulatory network inference, this study has several limitations. First, the single-cell CRISPRi perturbation dataset was derived exclusively from the MDA-MB-231 cell line, representing a specific mesenchymal/basal-like TNBC model (Wang et al., 2025). Consequently, the reconstructed regulatory architecture may not fully capture the molecular and epigenetic heterogeneity observed across different TNBC subtypes and individual patients. Second, the physical transcription factor binding priors were obtained from ENCODE Factorbook clustered ChIP-seq datasets, which aggregate evidence across multiple human cell types rather than exclusively representing TNBC-specific chromatin occupancy. This may introduce context-dependent discrepancies in the inferred TF–enhancer interactions. Third, although CRISPRi perturbation provides interventional evidence for enhancer-to-gene regulatory relationships under Pearl’s do-calculus, the upstream TF-to-enhancer interactions remain supported by observational chromatin occupancy data; direct genetic validation of upstream TF occupancy will require multiplexed TF knockouts. Fourth, single-cell Perturb-seq measurements are subject to guide-assignment uncertainty, potential off-target effects, and technical pseudoreplication arising from multiple cells sharing the same guide RNA. Although guide-level cell aggregation was employed to mitigate some of these concerns, residual experimental and analytical biases may remain. Finally, the predicted therapeutic combinations were evaluated through computational network disruption simulations rather than direct pharmacological experiments. Therefore, the predicted network inhibition percentages do not establish actual drug synergy, therapeutic efficacy, tolerability, or clinical benefit.

## Future Perspectives

Future investigations should extend this framework across multiple TNBC molecular subtypes and patient-derived organoid models to evaluate the reproducibility and generalizability of the identified regulatory networks. Incorporating TNBC-specific ChIP-seq or CUT&RUN datasets would further improve the biological context specificity of transcription factor–enhancer interactions. Additionally, multiplexed transcription factor perturbation experiments, combined with single-cell transcriptomic profiling, could provide direct experimental validation of upstream regulatory relationships and strengthen the causal interpretation of the complete tripartite network. Improved guide-assignment strategies, independent biological replicates, and systematic assessment of potential off-target effects would also enhance the reliability of future Perturb-seq analyses. Most importantly, the computationally prioritized therapeutic combinations, particularly BET bromodomain, p300 HAT, and HDAC inhibition, should be evaluated through in vitro pharmacological experiments across diverse TNBC cell lines using established drug synergy assessment methods, such as the Chou–Talalay combination index or Bliss independence. Subsequent in vivo xenograft studies would be necessary to investigate antitumor efficacy, toxicity, and therapeutic index. Ultimately, integrating experimentally validated regulatory networks with patient-specific molecular profiles may facilitate the development of more precise, mechanism-driven combination therapeutic strategies for TNBC.

## Conclusion

In this investigation, we addressed a foundational challenge in single-cell systems biology: inferring directed regulatory networks by integrating functional perturbation responses with physical chromatin occupancy priors. By formulating an evidence-weighted tripartite framework that couples large-scale single-cell CRISPRi enhancer perturbations (95,678 cells across 11,975 sgRNAs) with ENCODE ChIP-seq priors across 819 somatic mutation-associated enhancers, we mapped 6,366,349 candidate regulatory paths supporting 542,760 unique TF–target gene functional relationships in human Triple-Negative Breast Cancer.

Our findings demonstrate that multi-omics evidence integration substantially improves precision and reduces false-positive inflation relative to observational co-expression baselines, achieving robust discriminatory accuracy (AUROC = 0.8999, AUPRC = 0.8745, FDR = 9.87%) on held-out test enhancers compared to GENIE3 (AUPRC = 0.6323, FDR = 35.73%) and co-expression (AUPRC = 0.6814, FDR = 32.27%). Directed multigraph topological analysis and PageRank prioritization identified CTCF, POLR2A, EP300, JUND, and MYC as central transcriptional drivers anchoring the TNBC regulatory network. Explainable AI (SHAP TreeExplainer) characterized non-linear regulatory dependencies, revealing transactivation thresholds in AP-1/JUND circuits. In silico driver feature silencing correlated strongly with experimental single-cell CRISPRi fold changes (R² = 0.473, Pearson r = 0.688, p = 3.37 × 10⁻¹²² across N = 864 genes), and prioritized master drivers significantly stratified overall survival in multivariate Cox models across primary TNBC cohorts (METABRIC, n = 320; TCGA-BRCA, n = 160).

Finally, in silico therapeutic perturbation modeling identified a prioritized tri-modal epigenetic combination (combining BET bromodomain inhibition [OTX015/Birabresib] and p300 HAT inhibition [A-485/CCS1477] with class I HDAC inhibition [Entinostat/Vorinostat]) capable of disrupting 93% of downstream oncogenic regulatory edges in the modeled network. Collectively, this research establishes an open-science, computationally reproducible framework integrating functional perturbation genomics and physical binding priors, providing prioritized candidate drivers and combination hypotheses to guide experimental follow-up in aggressive triple-negative breast cancer.

## Data and Code Availability

All experimental datasets, processed matrices, causal regulatory networks, and analytical code supporting the findings of this study are publicly available to ensure complete transparency, reproducibility, and compliance with FAIR (Findable, Accessible, Interoperable, and Reusable) data standards.

Primary Experimental Datasets: The raw and processed single-cell CRISPRi perturbational sequencing datasets for human MDA-MB-231 triple-negative breast cancer cells are deposited in the NCBI Gene Expression Omnibus (GEO) under accession GSE224986 (Wang et al., 2025). Genomic coordinates for targeted somatic mutation enhancers and library annotations are accessible under GEO accession GSE223812 (Wang et al., 2025). Genome-wide transcription factor ChIP-seq physical binding clusters across 340 factors were retrieved from the ENCODE Factorbook / UCSC Genome Browser repository (encRegTfbsClustered). Independent clinical breast cancer validation cohorts are publicly accessible via the Genomic Data Commons / cBioPortal (TCGA-BRCA, clinically annotated Triple-Negative / Basal-like subset, n = 160) and European Genome-phenome Archive (METABRIC, basal-like subset, n = 320) (METABRIC Group et al., 2012; The Cancer Genome Atlas Network, 2012).

Processed Causal Networks and Supplementary Data: All derived high-throughput matrices and analytical resources are openly accessible via Zenodo and the Supporting Information:

i. S1 Data: Single-cell quality control thresholds, sequencing depth metrics, and cohort parameters across 95,678 cells (S1_Data.xlsx);
ii. S1 Table: Full genome-wide physical TF–enhancer prior interaction matrix comprising 15,059 binding events across 161 unique transcription factors (S1_Table.xlsx);
iii. S2 Data: Interventional enhancer perturbation response matrix encompassing 1,110,566 candidate functional regulatory edges across 819 targeted enhancers, permanently archived on Zenodo at https://doi.org/10.5281/zenodo.23897809 (Supplementary_Data_2_Perturbation_Responses.csv.gz);
iv. S3 Data: Complete reconstructed tripartite causal Gene Regulatory Network encompassing 6,366,350 high-confidence directed regulatory paths (TF → Enhancer → Target Gene), permanently archived on Zenodo at https://doi.org/10.5281/zenodo.23897809 (Supplementary_Data_3_Tripartite_Causal_GRN.csv.gz); and
v. S1 Display: Standalone interactive 2D/3D force-directed physics graph of the causal network (S1_Display.html).

Computational Code Availability: The complete end-to-end Python computational pipeline is openly available on GitHub under the open-source MIT License at: https://github.com/Omarrfaroqueiu79/tnbc-causal-grn. The repository contains documented, reproducible scripts for sparse data ingestion, vectorized two-sample Welch t-testing (S1 Text [Supplementary Note 3]), tripartite network triangulation, Google PageRank scoring, TreeExplainer SHAP attribution modeling, in silico knockout simulations, and publication figure generation. All high-throughput perturbation response matrices and reconstructed causal network models are permanently archived and citable on Zenodo under DOI: 10.5281/zenodo.23897809 (https://doi.org/10.5281/zenodo.23897809). Study design, single-cell quality control, and clinical observational cohort modeling adhere to the STROBE reporting guidelines (S1 Checklist).

## Acknowledgments

The authors thank the ENCODE Consortium, the National Center for Biotechnology Information (NCBI) Gene Expression Omnibus, the Cancer Genome Atlas (TCGA), and the METABRIC consortium for maintaining open-access multi-omics and clinical repositories. Computational analyses were executed using high-performance scientific computing infrastructure at the Department of Statistics and Data Science, Islamic University, Kushtia.

## Supporting Information

**S1 Text**. Supplementary Notes, Mathematical Formulations, and Extended Tables. Detailed derivation of two-sample Welch t-test vectorization, Directed Acyclic Graph (DAG) identifiability under Pearl’s do-calculus, Google PageRank personalization vectors, TreeExplainer SHAP background distributions, and biochemical mechanisms of synergistic epigenetic combinations. Includes embedded Supplementary Tables S1–S8 (S1_Text.docx).

**S1 Table**. Full Genome-Wide Physical TF–Enhancer Prior Interaction Matrix. Complete catalog of 15,059 physical binding events across 161 unique transcription factors mapped to 819 targeted super-enhancer coordinates (S1_Table.xlsx).

**S1 Data**. Single-Cell Quality Control Thresholds and Dataset Parameters. Distribution of unique molecular identifiers (UMIs), gene detection counts, mitochondrial fraction filtering, and cell-cycle state distributions across 95,678 MDA-MB-231 cells (S1_Data.xlsx).

**S2 Data**. Interventional Enhancer Perturbation Response Matrix. Full differential expression response matrix comprising 1,110,566 candidate functional edges across 819 targeted enhancers. Permanently archived and openly accessible on Zenodo at https://doi.org/10.5281/zenodo.23897809 (Supplementary_Data_2_Perturbation_Responses.csv.gz).

**S3 Data**. Reconstructed Tripartite Causal Gene Regulatory Network. High-confidence directed regulatory paths (TF → Enhancer → Target Gene) partitioned across 6,366,350 triangulated edges. Permanently archived and openly accessible on Zenodo at https://doi.org/10.5281/zenodo.23897809 (Supplementary_Data_3_Tripartite_Causal_GRN.csv.gz).

**S1 Display**. Standalone Interactive Force-Directed Physics Graph. Interactive 2D/3D visualization of the prioritized triple-negative breast cancer causal regulatory network (S1_Display.html).

**S1 Checklist**. STROBE Statement Reporting Checklist. Completed STROBE reporting checklist for observational cohort modeling and single-cell quality control (S1_Checklist.docx).

